# Joint model shows association of Mapuche genetic ancestry and longitudinal BMI with early menarche

**DOI:** 10.64898/2025.12.04.692408

**Authors:** Lucas Vicuña, Cristian Meza, Danilo Alvares, Veronica Mericq, Ana Pereira, Susana Eyheramendy

**Affiliations:** Departamento de Laboratorios Clínicos, Escuela de Medicina, Pontificia Universidad Católica de Chile, Santiago, Chile; INGEMAT-CIMFAV, Faculty of Engineering, Universidad de Valparaíso, Valparaíso, Chile; MRC Biostatistics Unit, University of Cambridge, Cambridge, UK; Institute of Maternal and Child Research, Faculty of Medicine, Universidad de Chile, Santiago, Chile; Institute of Nutrition and Food Technology, Universidad de Chile, Santiago, Chile; Faculty of Engineering and Science, Universidad Adolfo Ibañez, Santiago, Chile; Instituto Milenio Fundamentos de los Datos (IMFD), Santiago, Chile; Data Observatory (DO), Santiago, Chile

**Keywords:** Body mass index, early menarche, genetic ancestry, mixed models, joint modeling

## Abstract

The age at puberty onset varies greatly between individuals and ethnic populations, with significant health implications. Early menarche increases risk for breast cancer, cardiovascular disease, depression, behavioral disorders, diabetes, and all-cause mortality. While genetic factors and higher body mass index (BMI) have been associated with earlier pubertal timing in girls, the effect of Native American genetic ancestry on menarche timing is virtually unknown. We assessed how individual Mapuche Native American genetic ancestry proportions and BMI changes over time influence age at menarche in a cohort of admixed girls with mainly European and Mapuche ancestries. We developed a novel joint statistical model that links a Bernoulli random variable with longitudinal trajectories. The model captures individual-specific BMI trajectory dynamics through individual random effects, linking longitudinal BMI patterns directly to early menarche risk, incorporating the assessment of genetic ancestry effects as well. Two parameter estimation methods were developed, confirming the robustness of the model fit.

We found significant ancestry effects, with girls experiencing early menarche having higher mean Mapuche ancestry proportions (47.5% vs 45.1%, *p* = 0.010). Each 10% increase in Mapuche ancestry substantially increased early menarche risk (*γ*_1_ = 3.405, *p* = 0.010). Childhood BMI growth patterns strongly predicted early menarche: both BMI change rate (*α*_1_ = 1.189, *p* < 0.001) and growth acceleration (*α*_2_ = 9.080, *p* = 0.003) were significant predictors, while baseline BMI showed no association (*p* = 0.254). These results demonstrate that higher Mapuche ancestry and accelerating BMI trajectories independently increase early menarche likelihood.

Body mass index, early menarche, genetic ancestry, mixed models, joint modeling

## Introduction

Puberty is a complex developmental transition from childhood to reproductive maturity, marked by significant physiological and hormonal changes. Its timing varies widely across individuals, sexes, and populations. Girls typically begin puberty earlier than boys, with normal onset ranging from 8–12 years in girls and 9–14 years in boys [1]. Deviations from typical timing—either early or late—have been associated with increased risks for a range of adverse health outcomes, including cancer, cardiometabolic diseases, and neurocognitive disorders [2].

Both genetic and environmental factors contribute to pubertal timing. Twin and family studies estimate that genetic factors account for over half of the variation in age at menarche [3, 4]. Environmental influences such as nutrition, physical activity, psychosocial stress, and body composition also play critical roles, in part by modulating the hypothalamic gonadotropin-releasing hormone (GnRH) pulse generator [5]. Over recent decades, the age at pubertal onset has declined globally, with the average age at breast budding decreasing by approximately three months per decade [6, 7], and the age at menarche falling from 17 years in the 1800s to around 13 years in the mid-1900s [8, 9, 10].

Higher BMI and adiposity are consistently associated with earlier menarche [11, 12], though they do not fully explain its timing. Socioeconomic status, diet, physical activity, and psychosocial stressors also contribute [13, 14, 15]. Ethnic differences have been observed, with non-Hispanic Black girls reaching menarche earlier than other groups [16]. However, ethnicity is a categorical and socially defined variable, which may obscure underlying biological variation. In contrast, individual genetic ancestry proportions offer a continuous and biologically grounded measure of inter-population genetic differences [17, 18]. In particular, the effect of Native American genetic ancestry and menarche timing is virtually unknown.

In Chile, data from the Growth and Obesity Chilean Study (GOCS) show that girls reach thelarche at a mean age of 9.2 years, pubarche at 9.7 years, menarche at 11.9 years, and peak height velocity at 10.6 years [19]. While some studies have examined the interaction between ethnicity and BMI in relation to early menarche [20], the joint effect of individual genetic ancestry proportions and BMI trajectories has not been estimated using integrated modeling approaches. Such joint modeling is relevant because it can reveal whether genetic ancestry and individual BMI growth patterns act as independent risk factors for early menarche—relationships that may be obscured when these factors are analyzed separately. This represents a key gap in understanding pubertal development in admixed populations.

To address this, we propose a joint modeling approach that captures BMI trajectories and their association with early menarche, incorporating individual Mapuche ancestry proportions. Our model combines a quadratic mixed-effects specification for BMI with a logistic regression for early menarche, sharing random effects across submodels. We compare joint estimation via the stochastic approximation EM (SAEM) algorithm [21, 22] with a two-stage approach. While joint estimation offers integrated inference and improved statistical efficiency [23], it is computationally intensive. Two-stage estimation is faster but may introduce bias [24, 25]. We apply both methods to data from the GOCS cohort to evaluate their performance and provide new insights into ancestry-informed pubertal development.

## Methods

### Participants

The “Growth and Obesity Chilean Cohort Study” (GOCS) [26] includes children with singleton births only, gestational age between 37−42 weeks, birth weight of 2, 500 to 4, 500 g, and no physical or psychological conditions that could severely affect growth. Among the 1196 participants of the cohort, 943 (489 girls and 454 boys) met all inclusion criteria. Every 6 months, secondary sex characteristics were evaluated by a single trained dietitian, with permanent supervision from the pediatric endocrinologist. Every 4 − 6 months girls were interviewed by phone to determine the age of first menstrual bleeding. A questionnaire was applied to differentiate from other diagnosis such as vaginal infections or other genitourinary conditions. Children were born in 2002–2003, live in six low-middle income counties from the southeast area of Santiago, Chile, and belong to the same socioeconomic group. Details of the design, objectives, and recruitment strategies of the GOCS have been described elsewhere [27, 28].

## Genotyping

We used SNP array data from GOCS adolescents obtained in a previous study [29]. Individuals were genotyped with the Infinium ®Multi-Ethnic Global BeadChip (Illumina). We used Plink 1.9 [30] to exclude 18 samples with call rate <0.98 (18 samples), 10 samples with gender mismatch, and one sample from each pair of highly related individuals (IBS >0.2). We excluded variants with a minor allele frequency (MAF) < 0.01, and variants following at least one of the following conditions: have heterozygous genotypes on male X chromosome, call genotypes on the Y chromosome in females, have high heterozigosity (± 3 SD from the mean), have >5% missing genotype data, have duplicated physical positions (one variant was kept from each duplicate pair) and show significant deviations from Hardy-Weinberg equilibrium (*P* = 1 × 10^−6^). We removed A-T and C-G transversions to avoid inconsistencies with the reference human genome. After quality control filtering, we obtained the final data set of 421 girls and 774, 433 autosomal SNPs.

## Global ancestry inference

Global (i.e. individual) ancestry proportions of the participants were estimated with Admixture [31] in unsupervised mode. We used Yoruba (YRI, n=108) as the reference population for the African ancestry and Iberian Populations in Spain (IBS, n=107) for the European ancestry [32]. We used a merged dataset of 11 Mapuche [33] and 73 Aymara [34] as the reference Native American population panel. The latter two groups represent the main Native American sub-ancestries of admixed Chileans. Using K=3 the software clearly identified 3 ancestral groups corresponding to European, African and Native American (**Figure** 1). We used the Mapuche ancestry proportion as a covariate in this study, given that our previous analysis of this cohort [29] showed mean Mapuche ancestry of 43.8% compared to only 2.6% Aymara ancestry [29], making Mapuche the predominant Native American component.

**Figure 1.**
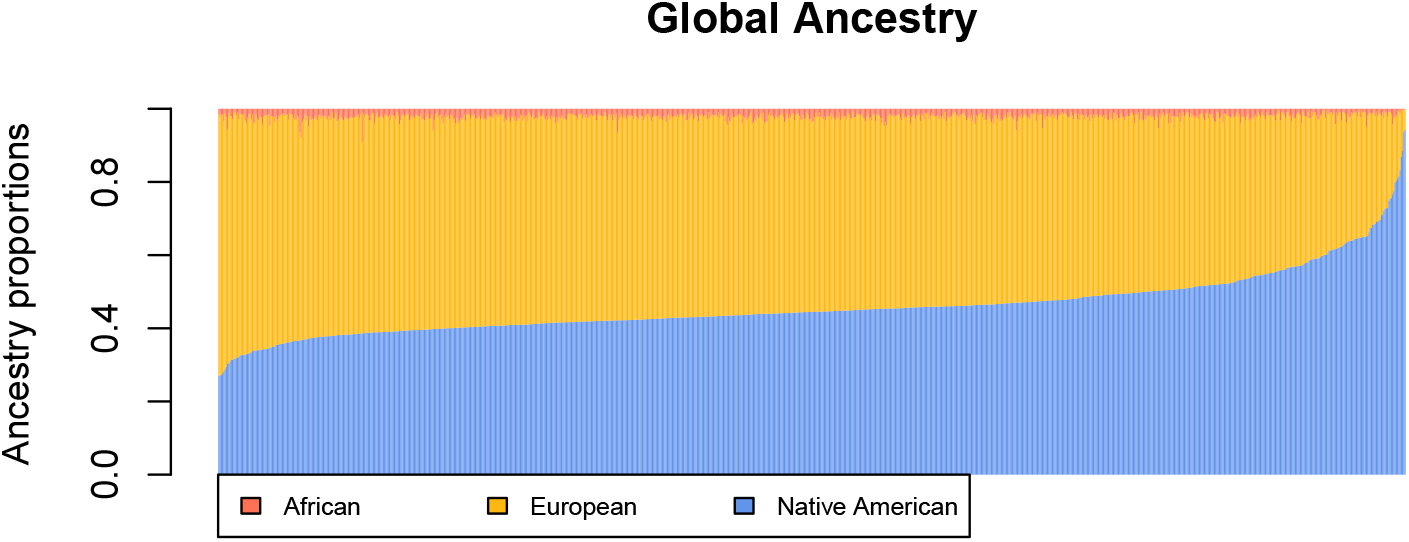
Individual ancestry proportions of the cohort. Each girl is represented by a vertical line.

### Earlier menarche threshold definition

Each population has its unique distribution that characterizes the age at menarche. It has been mentioned previously that, for example, Native American populations tend to have an earlier age at menarche compared to European populations. In this work we study the GOCS cohort, an admixed Chilean cohort. **Figure** 1 shows ancestry proportions of each girl in this study. To set a threshold that defines an earlier menarche we implemented a statistical model that empirically separated the ages in two groups. The model implemented is a mixture model [35] that assumes that the density of the ages can be generated from two Gaussian distributions. Specifically, we assume that the age at menarche of girl *i* follows a mixture of two Gaussians, 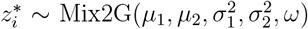, with its density function given by:

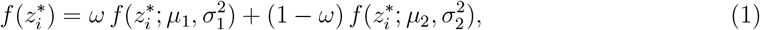

where *µ*_1_, *µ*_2_, 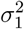, 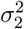 are the means and variances of the Gaussian densities *f*, where 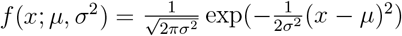, and 0 ≤ *ω* ≤ 1 denotes the weight for the first density. **Figure 2** (left panel) shows an histogram of the age at menarche, the estimation of the two Gaussian densities, and the earlier menarche threshold defined as the intersection point between the two densities. The first density (solid blue line) was estimated with 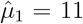 and 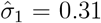, the second one (dashed green line) with 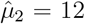 and 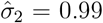. This estimated configuration led to an earlier menarche threshold at 11.5 years old. Therefore, we define *z*_*i*_ as a binary variable, where *z*_*i*_ = 1 if 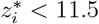 (an earlier menarche) and *z*_*i*_ = 0 otherwise.

**Figure 2.**
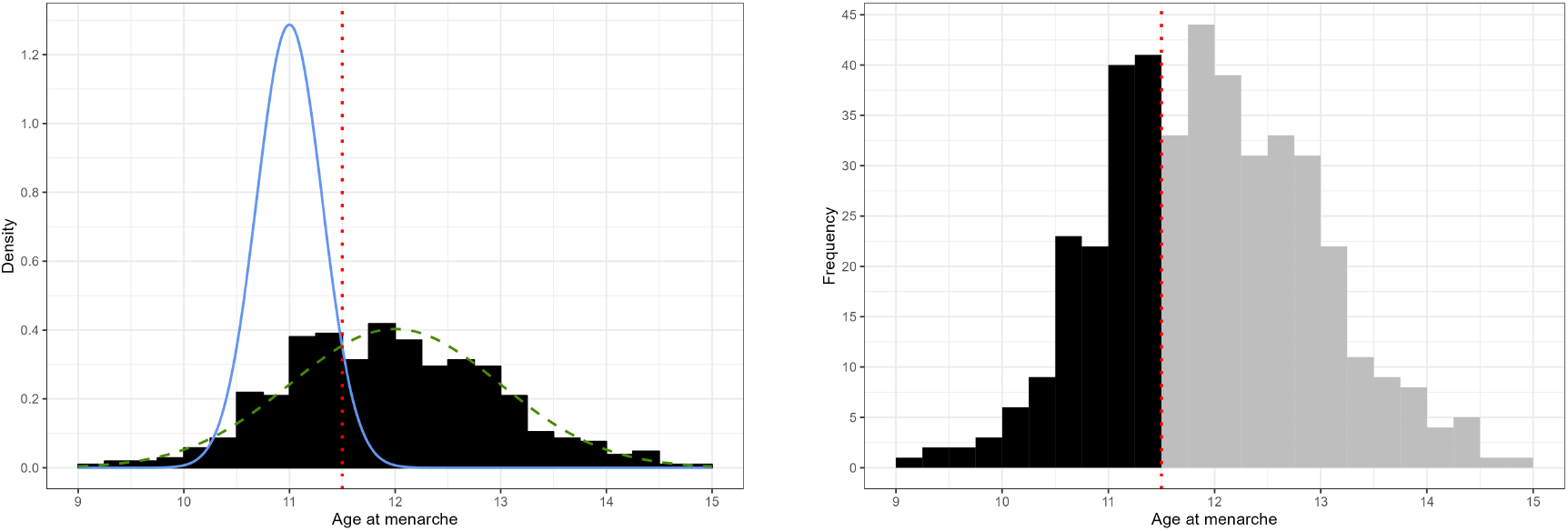
(Left) Distribution of the age at menarche (black bars), estimated densities for the Normal mixture (solid blue and dashed green lines), and early menarche threshold (dotted vertical red line). (Right) Distribution of the age at menarche. The dotted vertical red line indicates the earlier menarche threshold (11.5 years old) that defines the age at earlier (in black, *n* = 149) or later (in gray, *n* = 272) menarche.

**Figure** 3 shows the BMI longitudinal data for the two populations. In the left panel are the early menarche girls while at the right panel are the remaining girls.

**Figure 3.**
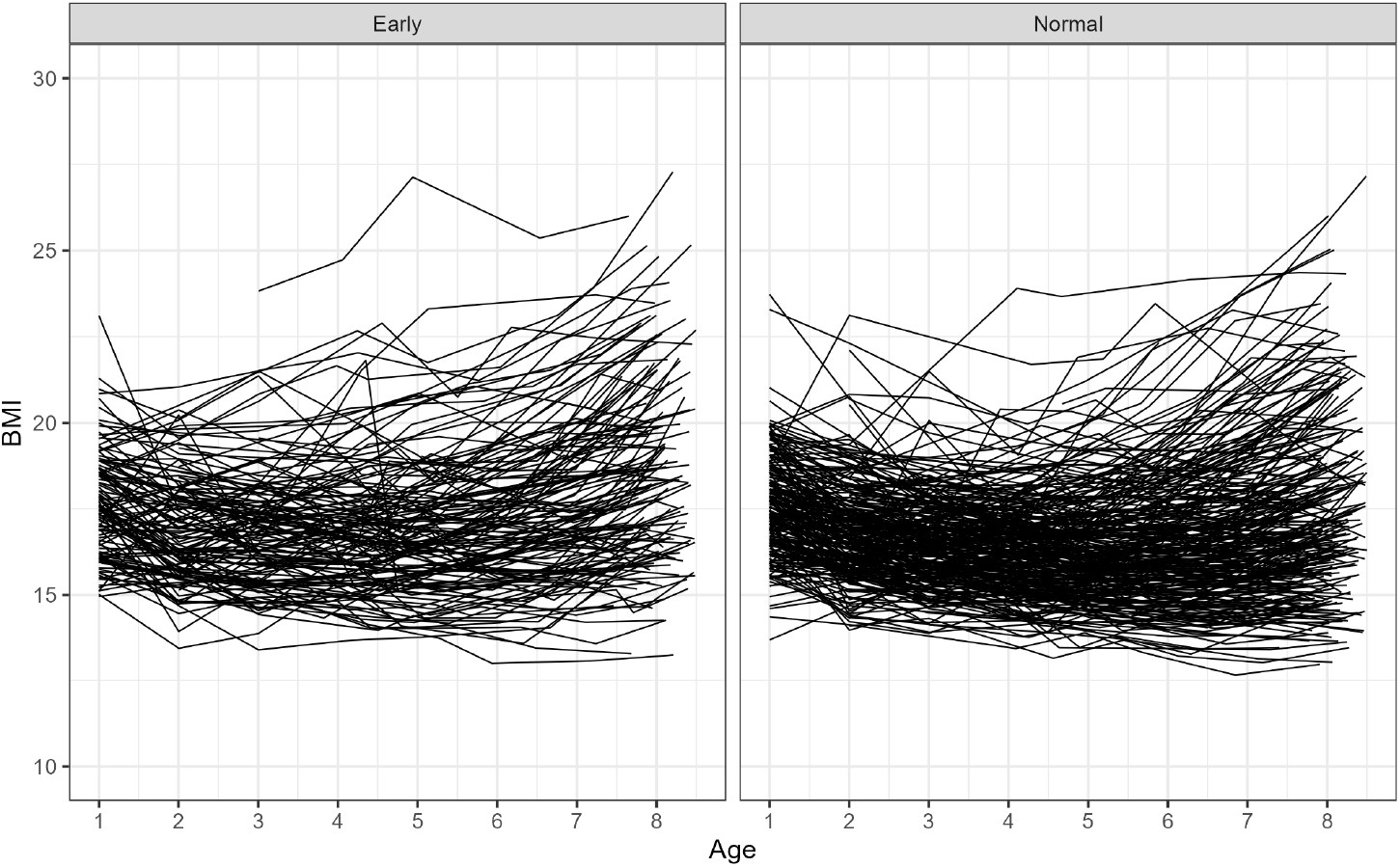
BMI versus age by menarche classification (Earlier: Age ≤ 11.5, Normal: Age > 11.5).

### Joint model

Our joint model proposal connects a longitudinal model to a logistic regression through random effects [36]. In particular, the longitudinal model describes the trajectory of the girls’ body mass index (BMI) over time while the logistic regression models the occurrence of early menarche. Specifically, let *y*_*ij*_ be the BMI associated with the *i*th girl at age *t*_*ij*_, for *i* = 1, …, *N* and *j* = 1, …, *n*_*i*_. We assume that BMI can be modeled by the following mixed-effects model [37]:

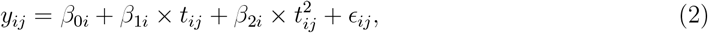

where the random effects ***β***_*i*_ = (*β*_0*i*_, *β*_1*i*_, *β*_2*i*_) follow a trivariate normal distribution with mean ***β*** = (*β*_0_, *β*_1_, *β*_2_) and variance-covariance matrix **∑**, and the error term is assumed ϵ_*ij*_ ∼ Normal(0, *σ*^2^).

Our primary response is defined as Normal (>11.5 years old) or Earlier (≤11.5 years old) menarche. From a modeling perspective, we can define *z*_*i*_ as a binary random variable, where 0 indicates that the *i*th girl’s age at menarche belongs to the Normal group and 1 otherwise. So, we propose to use a logistic regression model to explain the probability of early menarche, *p*_*i*_, as follows:

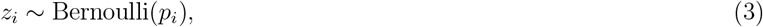

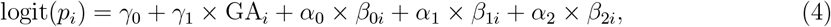

where GA_*i*_ represents the global Mapuche Native American ancestry proportion of the *i*th girl. Note that *β*_0*i*_, *β*_1*i*_ and *β*_2*i*_ are shared with the BMI submodel (2) and their respective coefficients, *α*_0_, *α*_1_ and *α*_2_, measure the strength of the association.

In the following subsections, we present two likelihood-based inference procedures for the joint model (2)-(3), the so-called *joint estimation*[38] and *two-stage estimation*.[39]

### Joint estimation via SAEM

The log-likelihood of the joint model (2)-(3) with parameters ***θ*** = (*β*_0_, *β*_1_, *β*_2_, *γ*_0_, *γ*_1_, *α*_0_, *α*_1_, *α*_2_, *σ*, **∑**) and random effects ***β*** = (***β***_1_, ***β***_2_, …, ***β***_*N*_) is given by:

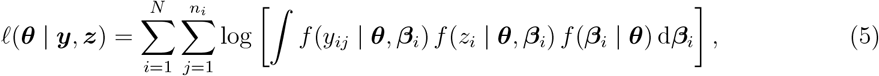

where 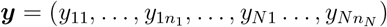, ***z*** = (*z*_1_, …, *z*_*N*_), and

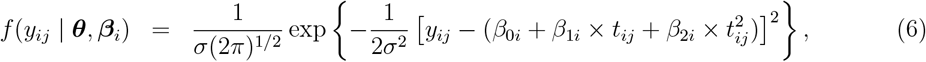

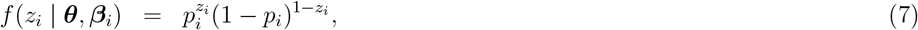

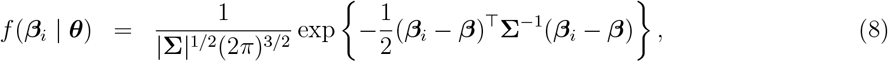

where *p*_*i*_ = 1/{ 1 + exp [− (*γ*_0_ + *γ*_1_ × GA_*i*_ + *α*_0_ × *β*_0*i*_ + *α*_1_ × *β*_1*i*_ + *α*_2_ × *β*_2*i*_)]}.

Treating the random effects ***β*** as non-observed (latent) data, we propose to maximize the log-likelihood described in (5) using the Stochastic Approximation (SA) version of the Expectation-Approximation (EM) algorithm, the so-called SAEM algorithm[21]. This algorithm is an iterative approach for the computation of maximum likelihood estimates in a wide variety of incomplete-data statistical problems. The SAEM algorithm consists of replacing the usual E-step of EM with a stochastic procedure. Algorithm 1 summarizes the procedure adopted by SAEM.

#### Algorithm 1

The SAEM algorithm

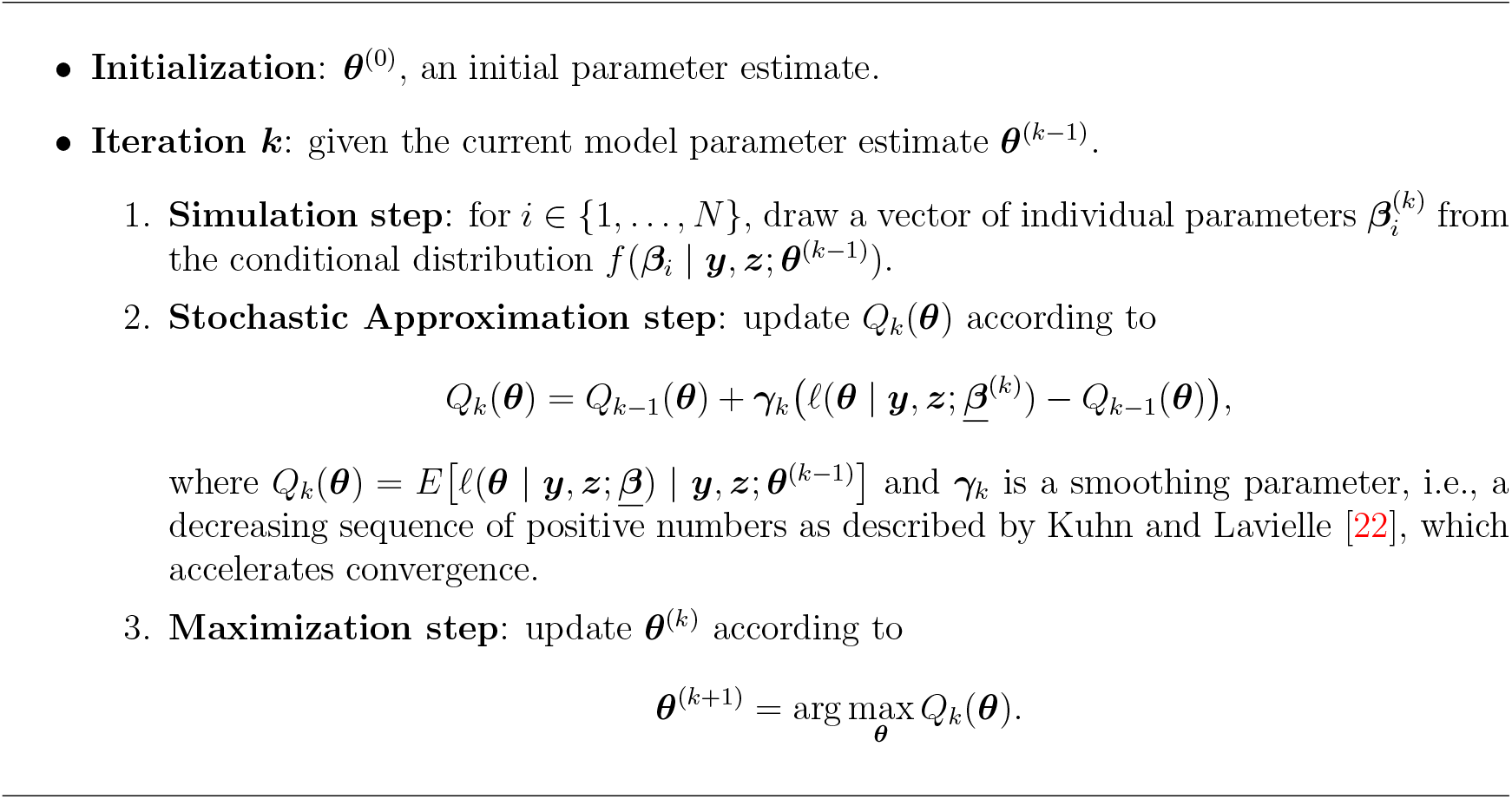

Kuhn and Lavielle [22] proposed to combine the SAEM with a Markov chain Monte Carlo (MCMC) procedure when the simulation step cannot be directly performed, as happens in joint models. Indeed, here, the conditional distribution of ***β*** cannot be computed in closed form. Then, the Simulation step in Algorithm 1 consists in applying the Metropolis-Hastings algorithm [40] with different kernels to approximate ***β***^(*k*)^ given ***y, z*** and ***θ***^(*k*−1)^.

SAEM technical and convergence details can be found in Delyon et a [21] and Kuhn and Lavielle [22] We used Monolix [41] software to implement the joint model (2)-(3) using SAEM estimation.

**Appendix C** of the Supplementary Material shows a guideline for using Monolix.

### Two-stage model

The inferential process of joint models is computationally demanding due to the presence of many random effects (which, from a frequentist perspective, must be integrated out) in a complex likelihood function. An appealing alternative is the so-called *two-stage estimation* [39]. This strategy proposes to fit each model separately, i.e., submodel (2) is fitted, and then the quantities of interest are extracted and used as known covariates to fit submodel (3). Mathematically, the first stage consists of estimating the shared information ***β***_*i*_ = (*β*_0*i*_, *β*_1*i*_, *β*_2*i*_) using the conditional distribution 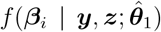, where 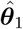 the maximum likelihood estimator of ***θ***_1_ = (*β*_0_, *β*_1_, *β*_2_, *σ*, **∑**) from the log-likelihood of the submodel (2):

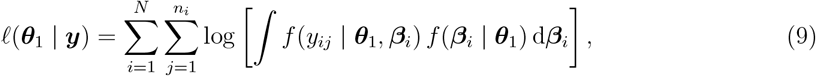

where *f* (*y*_*ij*_ | ***θ***_1_, ***β***_*i*_) and *f* (***β***_*i*_ | ***θ***_1_) are expressed as in (6) and (8), respectively. Then, in the second stage, the parameters ***θ***_2_ = (*γ*_0_, *γ*_1_, *α*_0_, *α*_1_, *α*_2_) are estimated from the log-likelihood of the submodel (3) assuming ***β***_1_, …, ***β***_*N*_ known (estimated values in the first stage):

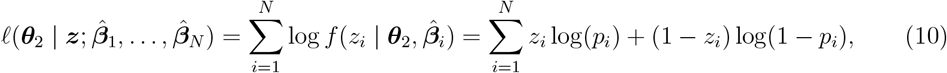

where 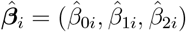 and 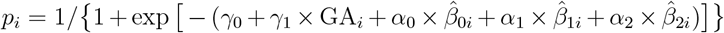. Although in some scenarios the two-stage approach produces slightly biased estimates [24, 25] it is widely used to reduce computational time in applications involving big data, such as genetics [42, 43] Another advantage of the two-stage estimation is its easy implementation using well-known functions in R language, such `lme` (nlme package) and `glm` (stats package).

## Results

Table 1 compares the results of the joint model (2)-(3) using both joint and two-stage estimations. The parameter estimates and their biological interpretations are equivalent between the two estimation approaches, as explained below.

**Table 1.** Estimated values of parameters, 95% confidence intervals, and p-values for joint and two-stage estimations using the joint model (2)-(3).

| Coeff | Joint Estimation (SAEM) |  |  | Two-stage Estimation |  |  |
| --- | --- | --- | --- | --- | --- | --- |
|  | Estimate | 95% CI | P-value | Estimate | 95% CI | P-value |
| $\beta_0$ | 18.184 | (18.014, 18.353) | <0.001 | 18.198 | (18.012, 18.384) | <0.001 |
| $\beta_1$ | -0.769 | (-0.835, -0.704) | <0.001 | -0.776 | (-0.851, -0.700) | <0.001 |
| $\beta_2$ | 0.092 | (0.085, 0.099) | <0.001 | 0.092 | (0.084, 0.100) | <0.001 |
| $\gamma_0$ | -3.875 | (-6.909, -0.841) | 0.016 | -3.758 | (-6.864, -0.695) | 0.016 |
| $\gamma_1$ | 3.405 | (0.822, 5.987) | 0.010 | 3.382 | (0.830, 6.015) | 0.010 |
| $\alpha_0$ | 0.097 | (-0.069, 0.263) | 0.254 | 0.091 | (-0.077, 0.259) | 0.286 |
| $\alpha_1$ | 1.189 | (0.482, 1.895) | <0.001 | 1.188 | (0.509, 1.884) | <0.001 |
| $\alpha_2$ | 9.080 | (3.059, 15.101) | 0.003 | 9.105 | (2.866, 15.454) | 0.004 |
| **-2log-like** | 8613.9 |  |  | 8603.4 |  |  |
| **AIC** | 8643.9 |  |  | 8633.4 |  |  |
| **BIC** | 8704.5 |  |  | 8712.7 |  |  |
| **Mean time (s)** | 153.6 |  |  | 3.5 |  |  |

### BMI trajectory parameters

The BMI trajectory parameters showed consistent estimates across methods: The baseline BMI at the reference age (*β*_0_ = 18.18, 95% CI: 18.01-18.35) represents the expected BMI when other factors are at their reference values. The linear age coefficient (*β*_1_ = −0.77, 95% CI: −0.84 to −0.70, *p* < 0.001) indicates an initial decrease in BMI with age, while the quadratic term (*β*_2_ = 0.092, 95% CI: 0.085-0.099, *p* < 0.001) captures the subsequent acceleration in BMI growth, reflecting the typical BMI trajectory during pubertal development.

### Genetic ancestry effects on the age at menarche

We inferred individual genetic ancestry proportions of the GOCS girls using Native American, European and African reference populations, as described in the Methods. **Figure** 1 shows the ancestry composition of our cohort, with Mapuche representing the predominant Native American component (mean 43.8%) compared to Aymara ancestry (2.6%) [29]. We found significant differences in mean Mapuche ancestry between the non-earlier group (Mean=0.451; SD=0.075) and the earlier group (Mean=0.475; SD=0.086). Girls with higher Mapuche ancestry showed increased risk of early menarche (*γ*_1_ = 3.41, 95% CI: 0.82-5.99, *p* = 0.010). This coefficient indicates that for every 10% increase in Mapuche ancestry proportion, the log-odds of early menarche increase by 0.34. The baseline risk parameter (*γ*_0_ = −3.88, 95% CI: −6.91 to −0.84, *p* = 0.016) establishes the reference probability of early menarche when Mapuche ancestry proportion equals zero and BMI trajectories are at their average values. These results suggest that, on average, girls with earlier menarche have higher Mapuche genetic ancestry proportions.

### BMI-menarche associations

Individual BMI patterns influence menarche timing in complex ways: The baseline BMI association (*α*_0_ = 0.097, 95% CI: −0.069 to 0.263, *p* = 0.254) was not statistically significant, suggesting that baseline BMI alone does not strongly predict early menarche. However, the linear BMI trajectory coefficient (*α*_1_ = 1.19, 95% CI: 0.48-1.90, *p <* 0.001) and quadratic trajectory coefficient (*α*_2_ = 9.08, 95% CI: 3.06-15.10, *p* = 0.003) were both significant, indicating that girls with steeper BMI increases over time have substantially higher odds of experiencing early menarche.

### Model performance metrics

The model performance metrics confirmed the equivalence of both approaches: The −2 log-likelihood values (8613.9 vs 8603.4), AIC scores (8643.9 vs 8633.4), and BIC values (8704.5 vs 8712.7) were very similar between joint and two-stage estimation. The primary trade-off was computational efficiency, with two-stage estimation requiring only 3.5 seconds compared to 153.6 seconds for joint estimation, while producing virtually identical biological conclusions.

Given the equivalence between the estimates detailed above, we validated our model using several diagnostic approaches. To analyze potential misspecifications in the structural model, **Figure** 4 (left panel) displays individual predictions versus observations as well as the 90% prediction interval. The low proportion of outliers (predictions outside the interval) and the symmetry between predictions and observations indicate good model fit and appropriate data modeling.

We also evaluated the performance of our BMI submodel using *visual predictive check* (VPC) plots [44]. This graphical tool compares different quantiles of the observed data to quantiles of simulated data (e.g., quantiles 10, 50, and 90). **Figure 4** (right panel) demonstrates that our longitudinal model has good predictive ability, particularly for the 10th and 90th percentiles, both for early and later ages.

**Figure 4.**
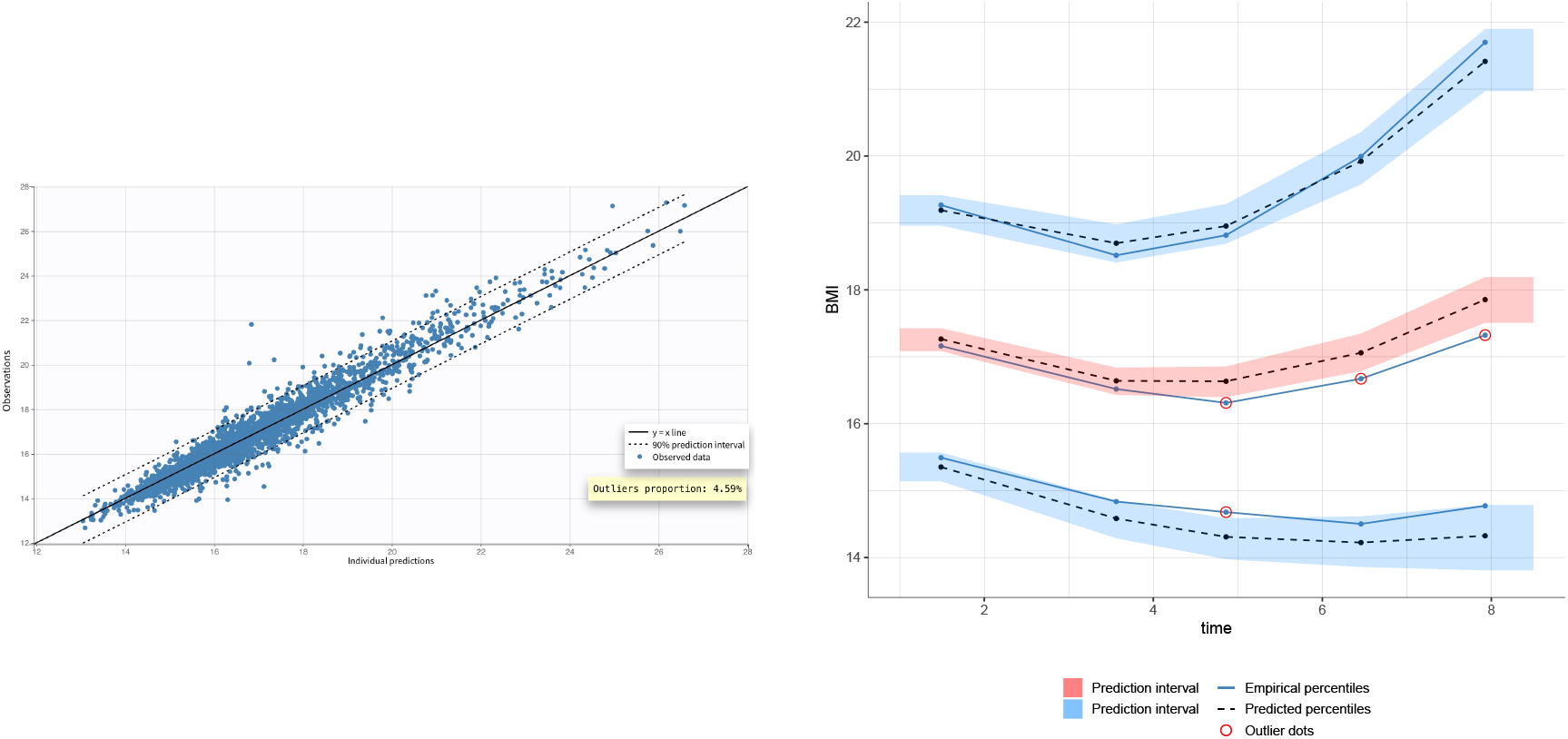
(Left) BMI observations vs predictions using the joint model (2)-(3) with SAEM estimation. (Right) Visual predictive check for BMI trajectories using the joint model (2)-(3) with SAEM estimation.

**SFigure** 1 in **Appendix A** of the Supplementary Material compares the distribution of the standardized random effects with standard Gaussian distributions. Since random effects should follow normal probability laws, this comparison validates our normality assumption.

In **Appendix B** of the Supplementary Material, we introduce a simpler alternative joint model where only two random effects (intercept and slope) are specified in the BMI submodel and shared with the early menarche submodel. This simpler longitudinal specification presents worse predictive performance (VPC and estimated random effects) as well as inferior goodness-of-fit metrics (likelihood, AIC, and BIC), supporting our choice of the quadratic specification.

## Discussion

This is the first study to show a positive association between Native American genetic ancestry and earlier age at menarche. Further, it provides the first quantitative assessment of how genetic ancestry proportions and BMI trajectories jointly influence early menarche timing in an admixed population.

Girls in the early menarche group had significantly higher mean Mapuche ancestry proportions (47.5% vs 45.1%), with an effect size (*γ*_1_ = 3.405, 95% CI: 0.822 − 5.987) indicating substantial ancestry-related risk. Each 10% increase in Mapuche ancestry increases the log-odds of early menarche by 0.34, independent of BMI growth patterns. This finding provides the first continuous, quantitative measure of Native American ancestry effects—specifically Mapuche ancestry—on age at menarche, extending beyond previous categorical ethnic comparisons [16, 45].

The joint modeling approach provides key methodological advantages over separate analyses by capturing individual-specific BMI trajectory dynamics through shared random effects. This reveals that personal growth patterns (linear and quadratic components) significantly predict early menarche (*α*_1_ = 1.189, p¡0.001; *α*_2_ = 9.080, p=0.003), while baseline BMI shows no association (*α*_0_ = 0.097, p=0.254)—a distinction masked in separate analyses using population-level measures. The shared random effects structure shows that genetic ancestry and BMI trajectories contribute independently and additively to early menarche risk, while providing integrated inference and improved statistical efficiency.

The association between early menarche and BMI has been extensively studied using linear regression [46, 47, 11], survival analysis [48, 49], and Mendelian randomization [50, 51]. However, the combined effects of genetic ancestry and longitudinal BMI trajectories have not been previously examined. Our joint modeling approach addresses this gap while incorporating continuous ancestry measures rather than categorical ethnicity classifications.

We compared joint estimation via the SAEM algorithm with two-stage approaches and found equivalent results, suggesting that two-stage estimation provides a computationally efficient alternative for this type of application. Our mathematically derived threshold for early menarche (11.5 years) using mixture modeling provides a population-appropriate cutoff for this admixed cohort, avoiding the limitations of fixed thresholds based on homogeneous populations.

For Chilean girls, those with higher Native American ancestry face elevated baseline risk, with additional risk from rapid BMI increases during childhood. Given established associations between early menarche and later health risks including breast cancer and cardiovascular disease, these findings support targeted interventions promoting healthy weight trajectories during childhood and adolescence, particularly in populations with higher indigenous ancestry.

We did not include socioeconomic status as a covariant in our models because the participants belong to the same socioeconomic group, live in the same six low-middle income counties, and a previous study on this same cohort demonstrated that children’s BMI was not affected by the maternal educational level, a proxy for socioeconomic status [52].

In conclusion, both individual genetic ancestry proportions and BMI growth trajectories independently influence early menarche timing in Chilean girls. These findings advance understanding of pubertal development in admixed populations with Native American and European ancestries and provide a framework for incorporating genetic ancestry into longitudinal health research, moving beyond categorical ethnic classifications toward more precise, biology-based approaches to population health disparities.

## Data availability

The code and simulation data used in this study are available at the following GitHub repository: https://github.com/lucas-vicuna/Menarche-2025

## Author contributions

S.E. conceived the study. C.M. and D.A. performed analyses and developed the algorithms. V.M. and A.P. performed anthropometric measurements and collected medical data. S.E., L.V., C.M. and D.A. wrote the manuscript and provided intellectual input. All the authors critically reviewed and accepted the final version.

## Ethical Approval

This study was approved by the Scientific Ethics Committee of Instituto de Nutrición y Tecnología en Alimentos (INTA).

## References

1. Rosenfield RL, Lipton RB and Drum ML. Thelarche, pubarche, and menarche attainment in children with normal and elevated body mass index. Pediatrics 2009; 123(1): 84–88.

2. Day FR, Elks CE, Murray A et al. Puberty timing associated with diabetes, cardiovascular disease and also diverse health outcomes in men and women: the UK Biobank study. Sci Rep 2015; 5: 11208.

3. Towne B, Czerwinski SA, Demerath EW et al. Heritability of age at menarche in girls from the fels longitudinal study. American Journal of Physical Anthropology 2005; 128(1): 210–219. DOI:10.1002/ajpa.20106. URL https://onlinelibrary.wiley.com/doi/abs/10.1002/ajpa.20106. https://onlinelibrary.wiley.com/doi/pdf/10.1002/ajpa.20106.

4. Dvornyk V and ul Haq W. Genetics of age at menarche: a systematic review. Human Reproduction Update 2012; 18(2): 198–210. DOI:10.1093/humupd/dmr050. URL https://doi.org/10.1093/humupd/dmr050. https://academic.oup.com/humupd/article-pdf/18/2/198/17055969/dmr050.pdf.

5. Herbison AE. Control of puberty onset and fertility by gonadotropin-releasing hormone neurons. Nature Reviews Endocrinology 2016; 12(8): 452–466. DOI:10.1038/nrendo.2016.70. URL http://dx.doi.org/10.1038/nrendo.2016.70.

6. Aksglaede L, rensen K, Petersen JH et al. Recent decline in age at breast development: the Copenhagen Puberty Study. Pediatrics 2009; 123(5): e932–939.

7. Eckert-Lind C, Busch AS, Petersen JH et al. Worldwide Secular Trends in Age at Pubertal Onset Assessed by Breast Development Among Girls: A Systematic Review and Meta-analysis. JAMA Pediatr 2020; 174(4): e195881.

8. Toppari J and Juul A. Trends in puberty timing in humans and environmental modifiers. Mol Cell Endocrinol 2010; 324(1-2): 39–44.

9. Sorensen K, Mouritsen A, Aksglaede L et al. Recent secular trends in pubertal timing: implications for evaluation and diagnosis of precocious puberty. Horm Res Paediatr 2012; 77(3): 137–145.

10. Wang Z, Asokan G, Onnela JP et al. Menarche and Time to Cycle Regularity Among Individuals Born Between 1950 and 2005 in the US. JAMA Network Open 2024; 7(5): e2412854–e2412854. DOI:10.1001/jamanetworkopen.2024.12854. URL https://doi.org/10.1001/jamanetworkopen.2024.12854. https://jamanetwork.com/journals/jamanetworkopen/articlepdf/2819141/wang_2024_oi_240446_1715884806.16605.pdf.

11. Biro FM, Pajak A, Wolff MS et al. Age of menarche in a longitudinal us cohort. J Pediatr Adolesc Gynecol 2018; 31(4): 339–345.

12. Rona R and Pereira G. Factors that influence age of menarche in girls in santiago, chile. Human Biology 1974; 46(1): 33–42. URL http://www.jstor.org/stable/41459926.

13. Cheng M, Yao Y, Zhao Y et al. The influence of socioeconomic status on menarcheal age among chinese school-age girls in tianjin, china. European Journal of Pediatrics 2021; 180(3): 825–832. DOI:10.1007/s00431-020-03803-4. URL https://doi.org/10.1007/s00431-020-03803-4.

14. Yermachenko A and Dvornyk V. Nongenetic determinants of age at menarche: A systematic review. BioMed Research International 2014; 2014(1): 371583. DOI:10.1155/2014/371583. URL https://onlinelibrary.wiley.com/doi/abs/10.1155/2014/371583. https://onlinelibrary.wiley.com/doi/pdf/10.1155/2014/371583.

15. Nguyen NTK, Fan HY, Tsai MC et al. Nutrient intake through childhood and early menarche onset in girls: Systematic review and meta-analysis. Nutrients 2020; 12(9). DOI:10.3390/nu12092544. URL https://www.mdpi.com/2072-6643/12/9/2544.

16. Chumlea WC, Schubert CM, Roche AF et al. Age at Menarche and Racial Comparisons in US Girls. Pediatrics 2003; 111(1): 110–113. DOI:10.1542/peds.111.1.110. URL https://doi.org/10.1542/peds.111.1.110. https://publications.aap.org/pediatrics/article-pdf/111/1/110/1006030/pe0103000110.pdf.

17. Bryc K, Auton A, Nelson MR et al. Genome-wide patterns of population structure and admixture in west africans and african americans. Proceedings of the National Academy of Sciences 2009; 107(2): 786–791. DOI:10.1073/pnas.0909559107. URL http://dx.doi.org/10.1073/pnas.0909559107.

18. Burchard EG, Ziv E, Coyle N et al. The importance of race and ethnic background in biomedical research and clinical practice. New England Journal of Medicine 2003; 348(12): 1170–1175. DOI:10.1056/nejmsb025007. URL http://dx.doi.org/10.1056/NEJMsb025007.

19. Pereira A, Corvalan C, Merino PM et al. Age at Pubertal Development in a Hispanic-Latina Female Population: Should the Definitions Be Revisited? J Pediatr Adolesc Gynecol 2019; 32(6): 579–583.

20. Kelly Y, Zilanawala A, Sacker A et al. Early puberty in 11-year-old girls: Millennium cohort study findings. Archives of Disease in Childhood 2017; 102(3): 232–237.

21. Delyon B, Lavielle M and Moulines E. Convergence of a stochastic approximation version of the EM algorithm. Annals of Statistics 1999; 27(1): 1028–1039.

22. Kuhn E and Lavielle M. Maximum likelihood estimation in nonlinear mixed effects models. Computational Statistics and Data Analysis 2005; 49(4): 1020–1038.

23. Alvares D, Armero C, Forte A et al. Sequential Monte Carlo methods in Bayesian joint models for longitudinal and time-to-event data. Statistical Modelling 2021; 21(1–2): 161–181.

24. Ibrahim JG, Chu H and Chen LM. Basic concepts and methods for joint models of longitudinal and survival data. Journal of Clinical Oncology 2010; 28(16): 2796–2801.

25. Zhang N, Chen H and Zou Y. A joint model of binary and longitudinal data with nonignorable missingness, with application to marital stress and late-life major depression in women. Journal of Applied Statistics 2014; 41(5): 1028–1039.

26. Pereira A, Garmendia ML, González D et al. Breast bud detection: a validation study in the chilean growth obesity cohort study. BMC Womens Health 2014; 14: 96. DOI:10.1186/1472-6874-14-96.

27. Kain J, Corvalán C, Lera L et al. Accelerated growth in early life and obesity in preschool chilean children. Obesity 2009; 17(8): 1603–1608.

28. Corvalan C, Uauy R, Kain J et al. Obesity indicators and cardiometabolic status in 4-y-old children. The American journal of clinical nutrition 2010; 91(1): 166–174.

29. Vicuna L, Norambuena T, Miranda JP et al. Novel loci and Mapuche genetic ancestry are associated with pubertal growth traits in Chilean boys. Hum Genet 2021; 140(12): 1651–1661.

30. Purcell S, Neale B, Todd-Brown K et al. Plink: a tool set for whole-genome association and population-based linkage analyses. Am J Hum Genet 2007; 81(3): 559–75. DOI:10.1086/519795.

31. Alexander DH, Novembre J and Lange K. Fast model-based estimation of ancestry in unrelated individuals. Genome Res 2009; 19(9): 1655–1664. DOI:10.1101/gr.094052.109. URL https://doi.org/10.1101/gr.094052.109.

32. 1000 Genomes Project Consortium, Auton A, Brooks LD et al. A global reference for human genetic variation. Nature 2015; 526(7571): 68–74. DOI:10.1038/nature15393.

33. Vidal EA, Moyano TC, Bustos BI et al. Whole genome sequence, variant discovery and annotation in Mapuche-Huilliche native south americans. Sci Rep 2019; 9(1): 2132.

34. Crawford JE, Amaru R, Song J et al. Natural selection on genes related to cardiovascular health in high-altitude adapted andeans. Am J Hum Genet 2017; 101(5): 752–767.

35. Frühwirth-Schnatter S, Celeux G and Robert CP. Handbook of mixture analysis. 1st ed. New York, NY, USA: Chapman & Hall/CRC, 2019.

36. Horrocks J and van Den Heuvel, M J. Prediction of pregnancy: a joint model for longitudinal and binary data. Bayesian Analysis 2009; 4(3): 523–538.

37. Laird LM and Ware JH. Random-effects models for longitudinal data. Biometrics 1982; 38(4): 963–974.

38. Rizopoulos D. Joint models for longitudinal and time-to-event data: with applications in R. 1st ed. Boca Raton, FL, USA: Chapman & Hall/CRC, 2012.

39. Tsiatis AA, DeGruttola V and Wulfsohn MS. Modeling the relationship of survival to longitudinal data measured with error. Applications to survival and CD4 counts in patients with AIDS. Journal of the American Statistical Association 1995; 90(429): 27–37.

40. Brooks S, Gelman A, Jones G et al. Handbook of Markov chain Monte Carlo. 1st ed. New York, NY, USA: Chapman & Hall/CRC, 2011.

41. Lixoft. Monolix version 2023r1. A Simulations Plus company, 2023. URL http://lixoft.com/products/monolix/.

42. Canouil M, Balkau B, Roussel R et al. Jointly modelling single nucleotide polymorphisms with longitudinal and time-to-event trait: An application to type 2 diabetes and fasting plasma glucose. Frontiers in Genetetics 2018; 9: 1–12.

43. Brossard M, Paterson AD, Espin-Garcia O et al. Characterization of direct and/or indirect genetic associations for multiple traits in longitudinal studies of disease progression. Genetics 2023; 225(1): 1–23.

44. Bergstrand M, Hooker AC, Wallin JE et al. Prediction-corrected visual predictive checks for diagnosing nonlinear mixed-effects models. The AAPS Journal 2011; 13(2): 143–151.

45. Ossa X, Bustos P, Muñoz S et al. Edad de menarquia y ascendencia indígena: Un estudio poblacional en chile. Revista médica de Chile 2012; 140(8): 1035–1042. DOI:10.4067/s0034-98872012000800010. URL http://dx.doi.org/10.4067/S0034-98872012000800010.

46. Askelund AD, Wootton RE, Torvik FA et al. Assessing causal links between age at menarche and adolescent mental health: a mendelian randomisation study. BMC Medicine 2024; 22(1): 155.

47. Asrullah M, L’Hoir M, Feskens EJM et al. Trend in age at menarche and its association with body weight, body mass index and non-communicable disease prevalence in indonesia: evidence from the indonesian family life survey (ifls). BMC Public Health 2022; 22(1): 628.

48. Lee JJ, Cook-Wiens G, Johnson BD et al. Age at menarche and risk of cardiovascular disease outcomes: Findings from the national heart lung and blood institute‐sponsored women’s ischemia syndrome evaluation. Journal of the American Heart Association 2019; 8(12): e012406.

49. Gil YJ, Park JH and Sung J. Discrete-time survival analysis of risk factors for early menarche in korean schoolgirls. J Prev Med Public Health 2023; 56(1): 59–66.

50. Xue P, Wang D, Chen Y et al. Association between body fat distribution and age at menarche: a two sample mendelian randomization study. Frontiers in Pediatrics 2024; 12.

51. Mumby HS, Elks CE, Li S et al. Mendelian Randomisation Study of Childhood BMI and Early Menarche. J Obes 2011; 2011: 180729.

52. Vicunã L, Barrientos E, Norambuena T et al. New insights from gwas on bmi-related growth traits in a longitudinal cohort of admixed children with native american and european ancestry. iScience 2023; 26(2): 106091. DOI:10.1016/j.isci.2023.106091. URL http://dx.doi.org/10.1016/j.isci.2023.106091.

